# Enteric glia adopt an activated pro-inflammatory state in response to human and bacterial amyloids

**DOI:** 10.1101/2022.08.08.503156

**Authors:** Peter Verstraelen, Samuel Van Remoortel, Nouchin De Loose, Rosanne Verboven, Gerardo Garcia-Diaz Barriga, Anne Christmann, Manuela Gries, Cagla Tükel, Sales Ibiza Martinez, Karl-Herbert Schäfer, Jean-Pierre Timmermans, Winnok H. De Vos

## Abstract

Mounting evidence suggests a role for the microbiome-gut-brain axis in amyloid-associated neurodegeneration, but the pathogenic changes induced by amyloids in the gastro-intestinal tract remain elusive. To scrutinize the early response to amyloids of human and bacterial origin, we challenged primary murine myenteric networks with Aβ_1-42_ (vs a scrambled version of Aβ_1-42_) and curli (vs culture medium), respectively, and performed shotgun RNA sequencing. Both amyloid types induced a transcriptional signature of DNA damage and cell cycle dysregulation. Using *in vitro* neurosphere-derived cultures and *in vivo* amyloid injections we found that enteric glia and smooth muscle cells were the most responsive cell types, showing increased proliferation, γH2AX burden and SOD2 levels after amyloid challenge. Consistent with this activated state, we identified a pro-inflammatory hub in the transcriptional profile of amyloid-stimulated myenteric networks. Enteric glia were the principal source of the associated cytokines, and *in vivo*, this was accompanied by an influx of immune cells. Together, these results shed new light on the intrinsic vulnerability of ENS cells to both amyloid species and position enteric glial cell activation as an early driver of neurodegenerative disease progression.

**Significance statement:** The increasing socio-economic impact of Alzheimer’s disease (AD), long sub-clinical disease progression window, and failure of drug candidates demand mechanistic insight into the early stages of disease development. Epidemiological associations and experimental studies in rodents suggest that the gut may be vulnerable to amyloids and mediate their transfer to the brain. However, whether and how amyloids induce local pathology in the gastro-intestinal wall is not known. We identified a pathogenic program that becomes activated in the gastro-intestinal tract after exposure to amyloid β and curli (the main bacterial amyloid), and show that enteric glia are responsible for creating an amyloid-induced pro-inflammatory environment. This insight of an early response in a distant, more accessible organ than the brain, may have important implications for both disease diagnosis and therapy.

## Introduction

Uptake of nutrients in the gastro-intestinal (GI) tract happens in an organ-autonomous fashion coordinated by the enteric nervous system (ENS), while the brain merely exerts modulatory functions. Recently, the gut-brain connection has gained attention since it might represent a more accessible and faster route for diagnosing and modulating sporadic neurodegenerative disorders like Alzheimer’s disease (AD)^1^. This was fueled by epidemiologic correlation between Inflammatory Bowel Disease (IBD, *i.e.,* recurrent GI inflammation) and dementia risk^2,3^, by parallel neuropathological manifestations in the brain and ENS of mouse models with a mutated Amyloid Precursor Protein (APP)^4,5^, and by experimental transfer of injected amyloids from the GI tract towards the brain in mice^6^.

An estimated 40% of bacterial species in our environment produce amyloidogenic proteins for cell attachment and biofilm formation^7–9^, many of which are also present in the GI tract. In addition, enteric neurons express amyloid precursor protein (APP)^10,11^, raising the probability for GI tissue to become exposed to amyloids from the outside and from within. It has been proposed that bacterial amyloids penetrate a leaky epithelial barrier during (inflamm)aging, that host-derived and human amyloids can seed each other’s aggregation, and that they use the same receptors (Toll Like Receptors, TLRs) to activate the innate immune system^12–14^. Curli, the principal amyloid produced by Gram-negative bacteria, was identified in a genome-wide screen as a bacterial product that promotes neurodegeneration in C. Elegans^15^, and experimental colonization with curli-producing bacteria promoted α-synuclein pathology in the GI tract and brain of mice and rats^16,17^. Similarly, injection of Aβ1-42 into the GI wall exacerbated amyloidosis in the n. vagus and brain of ICR mice^6,18^. In the AD brain, amyloid pathology is accompanied by chronic neuroinflammation, oxidative stress and accumulation of DNA damage^19–22^. However, studies on the local effects of amyloids in the ENS are lacking. Therefore, we have challenged myenteric networks with Aβ_1-42_ and curli, the archetypal human and bacterial amyloids^23^, and studied the downstream pathogenic events in different *in vitro* and *in vivo* models.

## Materials and methods

### Animal housing

Wild-type black6.N mice were bred and group-housed in the central animal facility at the University of Antwerp with food and water *ad libitum* and a dark/light cycle of 12/12h. All experimental procedures were approved by the ethical committee for animal testing of the University of Antwerp (file 2017-88).

### Amyloid preparation

Human amyloids Aβ_1-42_ (rPeptide A-1002-2), fluorescent Aβ_1-42_-hilyte555 (Anaspec AS-60480-01), scrambled Aβ_1-42_ (Aβ_scr_; rPeptide A-1004-1) or fluorescent Aβ_scr_-FAM (Anaspec AS-60892) were reconstituted in 1% NH_4_OH to obtain a 1 mM stock solution, bath sonicated and aliquoted for storage at −80 °C. The day before an experiment, an aliquot was dissolved in sterile PBS to a concentration of 10 μM, bath sonicated and allowed to oligomerize for 24h at 4 °C. Curli fibrils were isolated from Salmonella Typhimurium as previously described^24^. Mature curli fibrils (containing more nucleic acid) were used for the RNA sequencing, while intermediate fibrils were used for *in vitro* and injection experiments. These curli fibrils are devoid of lipopolysaccharide (LPS) and do not activate TLR4^25^. Fluorescent labeling of intermediate curli was performed using a HiLyte Fluor 555 protein labeling kit (Anaspec AS-72045). An aliquot of curli or curli-HiLyte555 was thawed from −80 °C, dissolved to the desired concentration, bath sonicated, and immediately used. For both amyloid types, low-binding Eppendorf tubes and filter tips were used to ascertain maximal recovery.

### Myenteric network isolation

Myenteric networks were isolated as described previously^26^. Briefly, animals were sacrificed via cervical dislocation and exsanguination, after which the entire colon was removed and transferred to a dish containing ice-cold dissection buffer (MEM with GlutaMAX and HEPES+ 1% PenStrep). The mesentery was removed and the colon cut open along the mesentery line. The muscularis externa was stripped off with fine forceps under a binocular microscope, cut into small (~25 mm^2^) pieces and enzymatically digested (0.4 U Liberase (Roche 5401151001) and 60 U DNAse I (Applichem A3778) in HBSS -Ca-Mg, 37 °C and 5% CO_2_ for 4.5h without shaking). Remaining smooth muscle cells were mechanically removed by gentle pipetting under a binocular stereomicroscope until the space between the ganglia was devoid of cells. The cleaned networks were transferred to a 48-well plate (4 wells per mouse, 3 mouse replicates, 12 wells in total) with 250 μl culture medium (DMEM-F12 with GlutaMAX (ThermoFisher 31331028), 2% B27 supplement (ThermoFisher 17504044), 1% bovine serum albumin, 0.1%β-mercaptoethanol, 1% PenStrep). After an overnight recovery period, the networks were stimulated with 1 μM oligomerized Aβ_1-42_ or Aβ_scr_, or an equivalent quantity mature curli fibrils (5 μg/ml) or non-supplemented DMEM-F12 medium. Exactly 24h after the amyloid addition, networks were lysed in RLT buffer + 1%β-mercaptoethanol for RNA isolation, and conditioned medium was collected for cytokine measurements via U-Plex Meso Scale Discovery analysis.

### mRNA sequencing

RNA was isolated using an RNeasy micro kit (Qiagen), its concentration measured with a Qubit device (ThermoFisher) and the integrity checked with a Bioanalyzer RNA pico chip (Agilent, RIN>8). cDNA libraries were prepared using a QuantSeq 3’ mRNA-Seq Library Prep kit FWD (Lexogen) and a qPCR add-on kit after which they were run on a Fragment Analyzer (Agilent), equimolar pooled and sequenced using an Illumina NextSeq 500/550 High Output Kit v2.5 (75 Cycles). Resulting reads were trimmed using the UrQt and SortMeRNA packages for R and aligned to the mouse reference genome (mm10) using Rsubread. Differentially expressed genes (DEGs) were identified using DESeq2 with standard settings (Benjamini-Hochberg-adjusted p-value cut-off at 0.1). Volcano plots were made in Graphpad Prism 9. Functional annotation, including GeneOntology term enrichment and construction of a network plot were done with Metascape using default settings^27^.

### Preparation of neurosphere-derived enteric glial/smooth muscle and neuronal cultures

Small and large intestines were dissected from E14 embryos and digested with 1 mg/ml DNAse 1 (AppliChem A3778) and 1 mg/ml collagenase A (Merck Millipore 10103586001) in DMEM-F12 at 37 °C while shaking. After 20 min, the partly digested intestines were pipetted up and down with a 100 μl pipette. After 45 min digestion, samples were filtered through a 70 μm cell strainer and cells were collected in DMEM-F12 medium supplemented with 1% glutamax, 1% HEPES, 1% sodium pyruvate and 1% PenStrep. They were centrifuged (5 min, 300g) and resuspended in DMEM-F12 medium additionally supplemented with 2% B27, 40 ng/ml EGF (ImmunoTools 12343407) and 20 ng/ml FGF (ImmunoTools 12343627). Cell material of 1 embryo was divided over 2 wells of a 6-well plate, in a volume of 2 ml per well. Neurospheres were allowed to grow for 1 week whereby growth factors were replenished at day 2 and an additional 1 ml of complete medium was added at day 4 after isolation. For final plating, supernatant containing non-attached neurospheres was collected and centrifuged (5 min, 300g). Attached neurospheres were briefly trypsinized, added to the same tube, and again centrifuged (5 min, 300g). To obtain enriched enteric glia cultures, neurospheres were plated onto PDL-coated 24- or 96-well plates in DMEM containing 10% FBS, 1% glutamax, 1% HEPES, 1% sodium pyruvate and 1% PenStrep. Glial cultures were used for experiments 5-7 days after final plating. To obtain neuronal cultures, neurospheres were plated onto PDL-coated 24- or 96-well plates in Neurobasal medium with 2% B27, 40 ng/ml GDNF (R&D Systems 512-GF-010), 1% glutamax, 1% HEPES and 1% PenStrep. The neuronal network was allowed to grow for 7 days before experimental treatments were started. Human oligomeric amyloids were used at a final concentration of 1 μM, and an equivalent quantity of intermediate curli fibrils (5 μg/ml) was used to stimulate neurosphere-derived cultures. LPS from *E. Coli* (InvivoGen tlrl-3pelps) was used at a final concentration of 100 ng/ml, and EdU (ThermoFisher C10338) was added for 4h at a final concentration of 10 μM and developed according to the manufacturer’s instructions.

### Intramural injections and whole mount preparation

Mice were injected at the age of 8 weeks. Anesthesia was induced with 5 and maintained with 2.5%isoflurane in O_2_. Animals were shaved and the abdomen washed with germicidal soap. The eyes were covered with an ophthalmologic gel and the animals were placed on a heating pad and covered with a sterile operation cloth. A pre-emptive subcutaneous injection with 0.05 mg/kg buprenorphine was administered, after which the abdominal cavity was opened along the *linea alba*. The caecum was exteriorized and regularly wetted with physiological solution. Oligomeric Aβ_1-42_-HiLyte555, Aβ_scr_, Aβ_scr_-FAM, fibrillar curli, fibrillar curli-HiLyte555 or sterile PBS was injected into the colon wall at 5 injection sites in a 1 cm region of the proximal colon. A total amount of 8 μg in 5 × 2 μl was injected using a 35G NanoFill needle. The region where the injections were given was marked by 2 final injections with tattoo ink. The abdominal muscles were fully closed by using continuous suture with 5.0 resolvable thread. The skin was then closed using subcutaneous sutures with 5.0 silk thread. Finally, a subcutaneous injection with 0.05 mg/kg buprenorphine was given before animals were placed under a heating lamp for recovery. The animals were placed in separate cages and closely monitored. The next morning, a final subcutaneous injection with 0.05 mg/kg buprenorphine was given. Mice were sacrificed by cervical dislocation, exactly 2 hours, 3 or 7 days after intramural injection. The proximal colon was dissected out and flushed with ice-cold Krebs solution. After removing the mesentery, the colon was opened along the mesentery line and pinned open in a black Sylgard Petri dish and fixed with 4% paraformaldehyde (PFA; 2h at room temperature (RT) for immunostaining or 24h at 4 °C for fluorescence *in situ* hybridization). Myenteric whole mounts were prepared by separating the external muscle layer from the submucosa/mucosa and removing the circular muscle layer under a binocular stereomicroscope.

### Immunostaining & microscopic imaging

All immunostaining steps were done in 96-well plates (50 μl/well) for cell cultures and in 1.5 ml Eppendorf tubes (150 μl/tube) at RT whilst gently shaking for whole mounts. Permeabilization was done in blocking buffer (0.1% bovine serum albumin, 10% normal horse serum (Innovative Research IGHSSER) in PBS) with 1% Triton X-100, for 5 min (cell cultures) or 2h (whole mounts). Primary antibodies (**Table 1**) were applied in blocking buffer for 4h (cultures) or 48h (whole mounts), followed by a PBS wash. Secondary antibodies (**Table 1**) were applied 2h (cell cultures) or overnight (whole mounts), followed by 10 min incubation with 4’,6-diamidino-2-phenylindole (DAPI, 2.5 μg/ml) and a final PBS wash. For cell cultures, HCS CellMask Deep Red Stain (ThermoFisher H32721, 2 μg/ml) was added along with secondary antibodies. Whole mounts were cover slipped in Citifluor (EMS 17970-100) with the side of the myenteric plexus facing the cover glass. Multichannel Z-stacks were acquired on a spinning disk confocal microscope (Ultra*VIEW* VoX, PerkinElmer) with 20X air and 60X oil immersion objectives (NA 0.75 and 1.4, respectively), or on a Nikon CSU-W1-SoRa spinning disk system with a 100X silicone immersion objective (NA 1.35). To obtain overview images of injection sites, 3×3 tiles were recorded with 10% overlap followed by flatfield correction and stitching in Fiji freeware^28^. Segmentation and quantification of cell images (nuclei, cells and γH2AX spots) were done with the in-house developed Fiji script CellBlocks (https://github.com/DeVosLab).

**Table 1.**
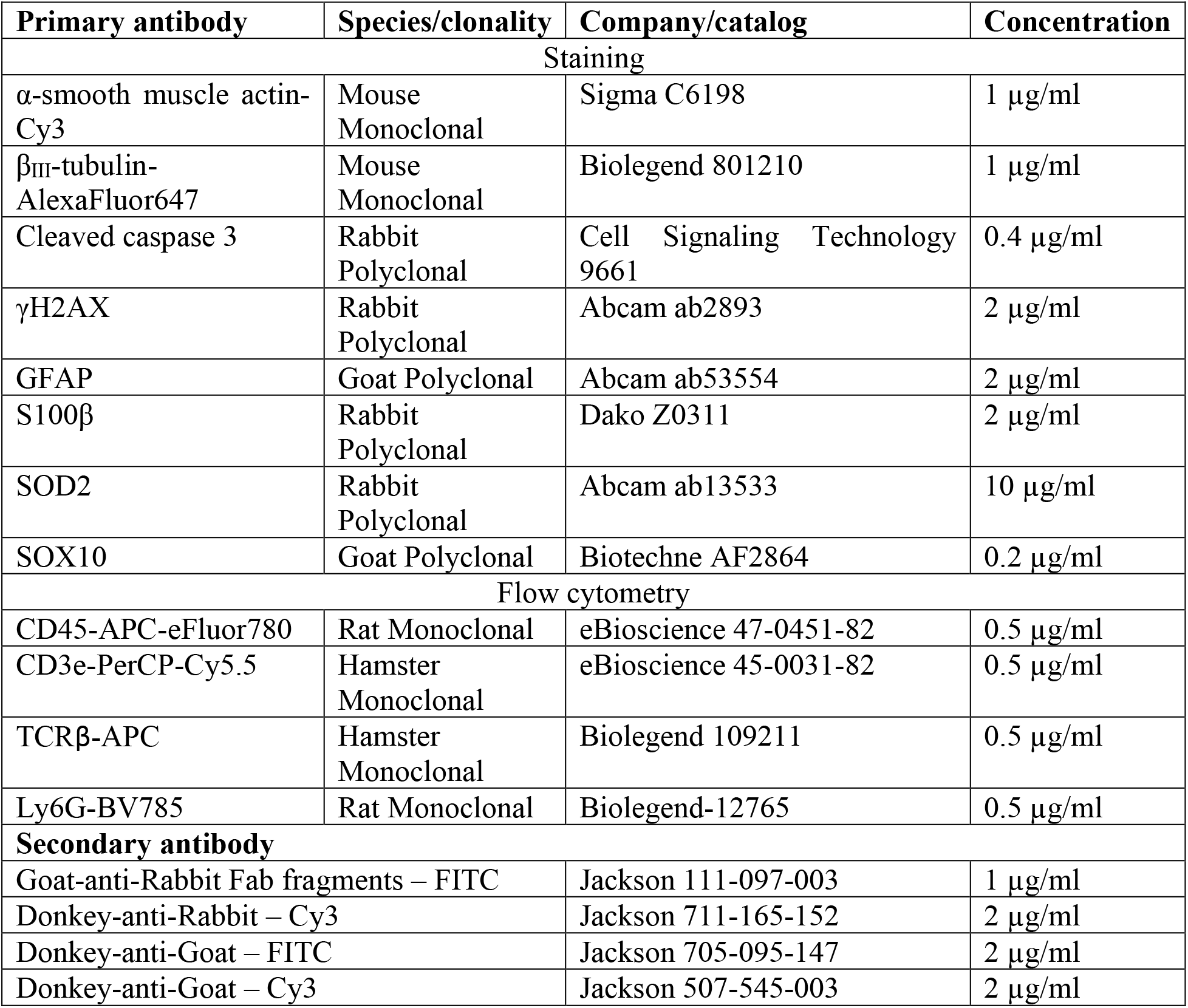
Primary and secondary antibodies

| Primary antibody | Species/clonality | Company/catalog | Concentration |
| --- | --- | --- | --- |
| Staining |  |  |  |
| α-smooth muscle actin-Cy3 | Mouse Monoclonal | Sigma C6198 | 1 μg/ml |
| β<sub>III</sub>-tubulin-AlexaFluor647 | Mouse Monoclonal | Biolegend 801210 | 1 μg/ml |
| Cleaved caspase 3 | Rabbit Polyclonal | Cell Signaling Technology 9661 | 0.4 μg/ml |
| γH2AX | Rabbit Polyclonal | Abcam ab2893 | 2 μg/ml |
| GFAP | Goat Polyclonal | Abcam ab53554 | 2 μg/ml |
| S100β | Rabbit Polyclonal | Dako Z0311 | 2 μg/ml |
| SOD2 | Rabbit Polyclonal | Abcam ab13533 | 10 μg/ml |
| SOX10 | Goat Polyclonal | Biotechne AF2864 | 0.2 μg/ml |
| Flow cytometry |  |  |  |
| CD45-APC-eFluor780 | Rat Monoclonal | eBioscience 47-0451-82 | 0.5 μg/ml |
| CD3e-PerCP-Cy5.5 | Hamster Monoclonal | eBioscience 45-0031-82 | 0.5 μg/ml |
| TCRβ-APC | Hamster Monoclonal | Biolegend 109211 | 0.5 μg/ml |
| Ly6G-BV785 | Rat Monoclonal | Biolegend-12765 | 0.5 μg/ml |
| Secondary antibody |  |  |  |
| Goat-anti-Rabbit Fab fragments – FITC | | Jackson 111-097-003 | 1 μg/ml |
| Donkey-anti-Rabbit – Cy3 | | Jackson 711-165-152 | 2 μg/ml |
| Donkey-anti-Goat – FITC | | Jackson 705-095-147 | 2 μg/ml |
| Donkey-anti-Goat – Cy3 | | Jackson 507-545-003 | 2 μg/ml |

### Quantitative PCR

Cells were lysed (RLT buffer with 1% β-mercaptoethanol) and RNA was isolated via column purification (NucleoSpin RNA kit, Macherey-Nagel 740955). RNA integrity and concentration were determined with BioAnalyzer (Agilent) and Nanodrop systems (Thermo Scientific), and 500 ng RNA (RIN>8) was transcribed to cDNA using the iScript first strand cDNA synthesis kit (Bio-Rad 1708891). qPCR was carried out on a 384-well Quantstudio Flex system (ThermoFisher) using the SsoAdvanced Universal SYBR Green master mix (Bio-Rad 1725272) with 0.5 μM forward and reverse primers (**Table 2**) and a 1:10 dilution of the cDNA. The protocol comprised an initial 30s denaturation step (95 °C), followed by 40 cycles of 10s at 95 °C and 30s at 60 °C, and finally a melting curve. Expression data was normalized to the reference genes, log2 transformed and imported into Genesis^29^, where it was normalized per gene before exporting the heatmap.

**Table 2.**
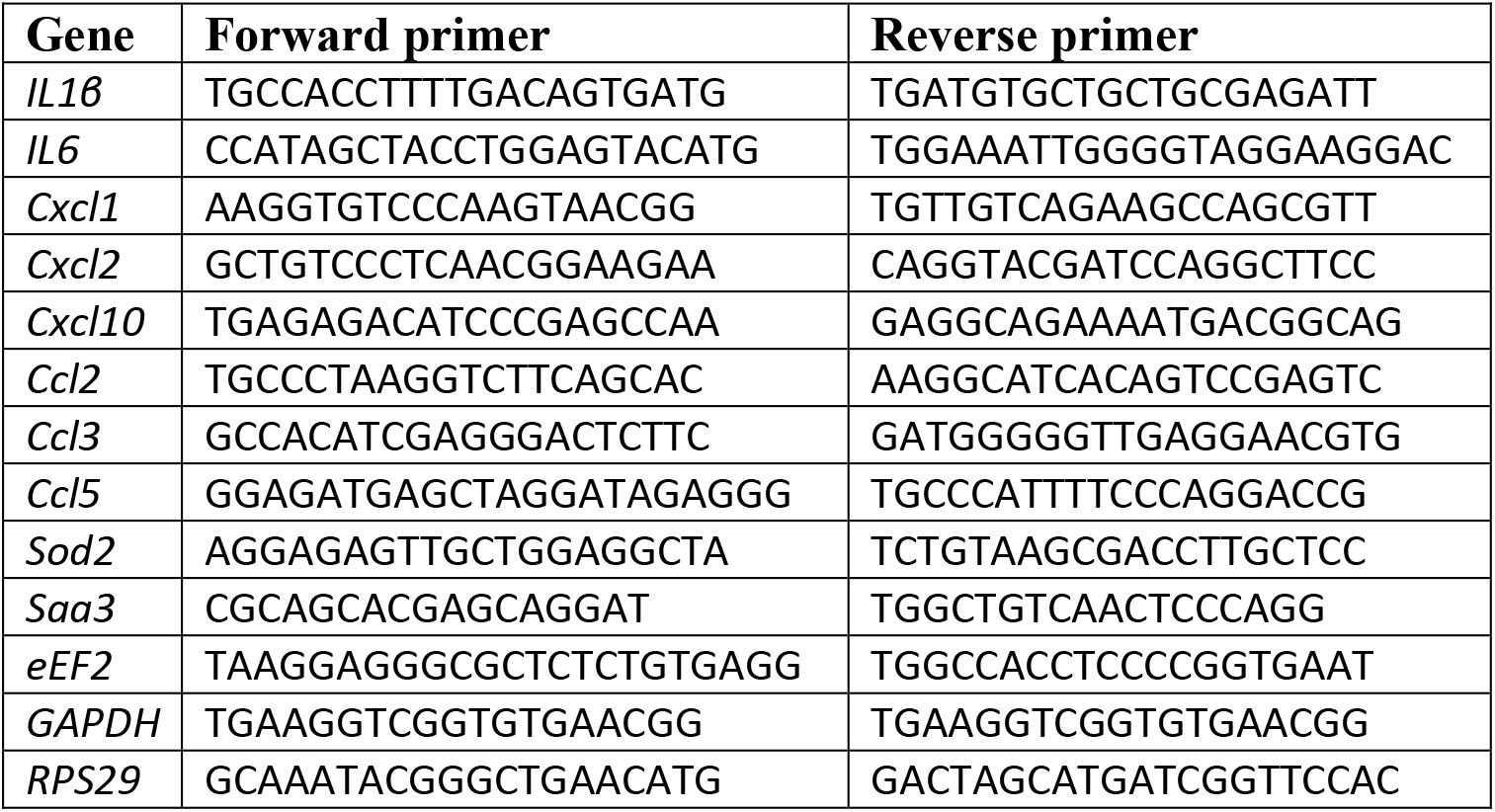
Primers for qPCR

### Fluorescence in situ hybridization

Fluorescence *in situ* hybridization analysis on whole mounts of the colonic myenteric plexus was performed using the Advanced Cell Diagnostics RNAscope Fluorescent Multiplex Kit (ACD 320850) according to the manufacturer’s instructions. After 24h fixation with 4% PFA at 4 °C, whole mounts were prepared and dehydrated through a graded ethanol series, followed by RNAscope Protease III treatment for 40 min at 40 °C. Tissue was then incubated overnight with fluorescent probe (RNAscope probe 437581 Mm-*Cxcl2*) at 40 °C under orbital shaking. After hybridization, whole mounts were washed twice with wash buffer and then processed for sequential hybridization using amplifier DNA (Amp1-FL, Amp 2-FL and Amp 3-FL) and fluorophore (Amp 4 Alt A-FL) at 40 °C for 30 min, 15 min, 30 min and 15 min, respectively. After hybridization, tissues were counterstained with SOX10 and DAPI, and mounted with ProLong Gold antifade reagent (Thermo Fisher P10144).

### Flow cytometry

A 3 cm piece of the proximal colon, containing the 1 cm injected region, was isolated 7 days after injection and the muscularis was removed with fine forceps under a binocular stereomicroscope. Small tissue pieces were digested with 0.5 mg/ml collagenase D (Roche 11088882001) and 5 U/ml DNAse 1 (Sigma-Aldrich 10104159001) in RPMI-1640 supplemented with 2% HEPES and 2% FBS for 30 min at 37 °C with continuous shaking. The resulting cell suspension was blocked using FACS buffer and passed through a 70 μm cell strainer, after which cells were centrifuged at 400g for 8 min at 4°C. Surface staining was performed by incubating cell suspensions for 20 min at 4°C with a mix of fluorescently conjugated antibodies (**Table 1**) in FACS buffer, followed by a PBS wash. To distinguish the live and dead cells, the cell pellets were resuspended in the Live/Dead Fixable Aqua Dead Cell Strain kit solution (ThermoFisher L34965) and incubated in the dark at RT for 30 min. Then, the cell pellets were fixed with 2% PFA at RT for 10 min, followed by a PBS wash and resuspension in FACS buffer. The samples were acquired using a BD FACSAria II Cell Sorter, and the obtained data were analyzed using FlowJo software (version 4.6.2, Treestar).

### Experimental design

RNA-Seq was performed on primary myenteric networks that were isolated from 3 mice and were divided over 4 wells per mouse (for Aβ_1-42_, Aβ_scr_, curli or medium treatment). Similarly, cytokine release was measured in myenteric networks from 8 mice, prepared on 2 different days and including the 3 mice that were used for RNA-Seq. Microscopic analyses on neurosphere-derived cultures were carried out on 6 wells with 12 images per well, originating from 3 independent cultures, *i.e.,* week-separated neurosphere-derived cultures prepared from different mothers (E14 embryos were pooled). Likewise, qPCR was carried out on 3 independent cultures. For all *in vivo* injections, 3 mouse replicates were considered for each treatment and time point, whereby 2 whole mounts could be prepared per mouse for different stainings. The number of Aβ^+^ myenteric neurons was assessed by manually counting 10 fields at 20X magnification per whole mount (amounting to a total of ~1.3 mm^2^/whole mount). The percentage of *Cxcl2^+^* glial nuclei was manually counted by an observer that was blinded for the treatment and normalized to the total number of SOX10^+^ glial nuclei in an image. At least 10 images per whole mount were considered which had on average 35 glial nuclei per image. Flow cytometry was performed on 4 mouse replicates per treatment type. Graphing and statistical analyses were carried out in Graphpad Prism 9 and SAS JMP Pro 14. The number of replicates, and results of statistical analyses are reported in the figure captions. Schematic representations of experimental protocols were created with BioRender.

## Results

### Human and bacterial amyloids trigger unique and shared transcriptional responses in primary myenteric networks

To determine whether amyloids affect the ENS, we isolated primary myenteric networks from WT Black6 mice and challenged them with human (Aβ_1-42_ vs Aβ_scr_) and bacterial (curli vs medium) amyloids. 24h later, we analyzed the transcriptome via bulk RNA-Seq (**Fig. 1A**). Inspection of typical cell type marker genes revealed that the primary networks consisted mainly of neurons, glia and smooth muscle cells, while immune cells were absent (**Fig. 1B**). Both amyloid types induced a transcriptional response whereby Aβ_1-42_ more potently disturbed myenteric network homeostasis (772 DEGs) than curli (228 DEGs; 53 shared; **Fig. 1C**), and this with good reproducibility between samples (**Fig. 1D**). This resulted in higher significance of the enriched gene ontology (GO) terms, with unique pathways such as glycerolipid metabolism being affected by Aβ_1-42_ but not by curli (-Log10 p-value, **Suppl. Fig. 1**). Being more interested in the shared response, we found that both amyloid types elicited a transcriptional hub of cell cycle dysregulation (**Fig. 1E**). Well-known cell cycle regulators such as *Pcna, Trp53, Ccnb1, Cdc20* and *Cdk1* were downregulated, while *Mdm2* was up after amyloid challenge, suggesting a global cell cycle wavering. In contrast, both amyloid types were represented in a hub concerning ‘muscle cell proliferation’, suggesting cell type-dependent cell cycle dysregulation. A DNA damage response was triggered more strongly by curli, while mitochondria and protein kinase B (Akt) signaling were disturbed mainly by Aβ_1-42_ (**Fig. 1E**). This illustrates that host-derived and bacterial amyloids are not inert but trigger a distinct transcriptional pathogenic response in the ENS.

**Figure 1.**
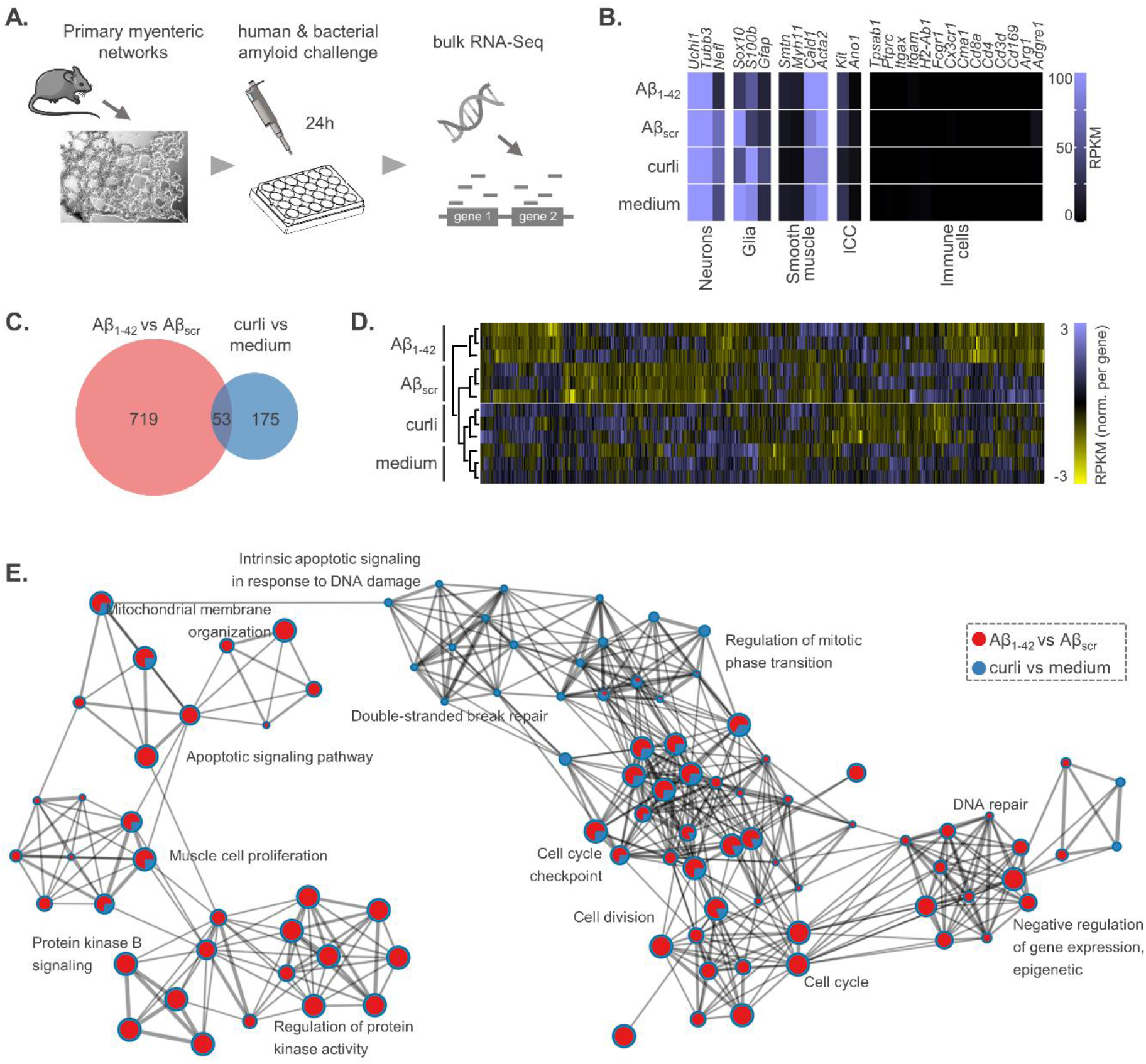
Human and bacterial amyloids trigger unique and shared transcriptional responses in primary myenteric networks. **A.** Experimental workflow. Primary myenteric networks were challenged with human (Aβ_1-42_ vs Aβ_scr_) or bacterial (curli vs PBS) amyloid for 24h and processed for bulk RNA-Seq (n=3 animals with 4 treatments/animal); **B.** Expression levels of typical cell type markers reveal that primary myenteric networks used for RNA-Seq predominantly consist of enteric neurons, glia and smooth muscle cells; **C.** Venn diagram depicting the number of DEGs in both comparisons; **D.** Heatmap of DEGs show that individual replicates of different treatments cluster and differentiate from the controls; **E.** Network plot of enriched GO terms represented as nodes, with node size proportional to the number of DEGs in the term and the color indicating the relative DEG counts of both treatments (shown as a pie chart).

### Enteric glia and smooth muscle cells become activated after amyloid challenge

Given that the cell cycle signature is unlikely to originate from the post-mitotic myenteric neurons, we challenged enteric neurosphere-derived glial cultures with human and bacterial amyloids (**Fig. 2A**). LPS, a well-known toxin of Gram-negative bacteria and TLR agonist, was included as a positive control since it has been described to induce astrocyte proliferation and DNA damage *in vitro*^30,31^. The cultures contain enteric glia (positive for GFAP and SOX10) but also smooth muscle cells (α-smooth muscle actin^+^ (αSMA); **Suppl. Fig. 2A**). Differential nuclei segmentation based on SOX10 or DAPI signal showed that both amyloid types increased EdU incorporation in glial as well as all (glia + smooth muscle) cells, indicating that both cell types adopt a proliferative state, almost up to similar levels as a treatment with LPS (**Fig. 2B**). Glial and smooth muscle cells also displayed an increased nuclear γH2AX spot occupancy, proxy for double-stranded DNA break repair, 72h after the amyloid addition (**Fig. 2C**). In correspondence with the transcriptomics data, we found that curli was a more potent inducer of a DNA damage response than Aβ_1-42_. We next asked whether amyloid-induced cell proliferation and DNA damage were associated with increased oxidative stress levels, as oxidative DNA damage was previously described in the AD brain^32^. We therefore immunostained challenged cultures for superoxide dismutase 2 (SOD2), a mitochondrial enzyme involved in reactive oxygen species clearance, which was also upregulated in the RNA-Seq dataset and found markedly higher SOD2 levels (**Fig. 2E**). γH2AX spot occupancy and SOD2 levels were also increased after LPS stimulation. Hence, our results suggest that enteric glia and smooth muscle cells adopt an activated state whereby they accumulate DNA damage, plausibly due to increased proliferation and oxidative stress.

**Figure 2.**
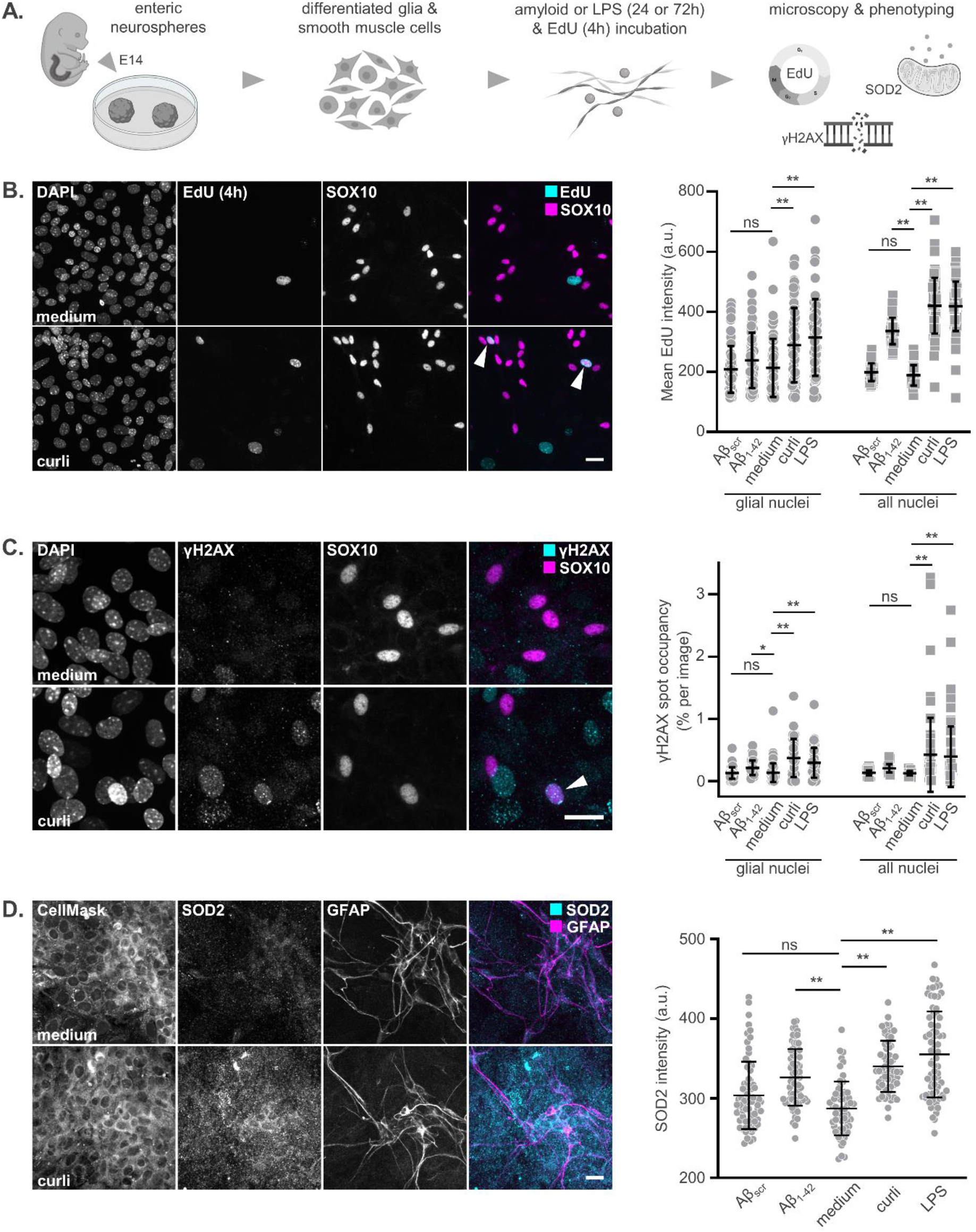
Enteric glia and smooth muscle cells become activated after amyloid challenge. **A.** Experimental workflow for challenging neurosphere-derived glial/smooth muscle cell cultures with amyloids and LPS and subsequent phenotyping (n=3 independent cultures with 24 images/culture); **B.** Representative images and EdU intensity quantification in glial nuclei as well as all nuclei (after resp. SOX10 and DAPI segmentation; arrowheads indicate EdU^+^SOX10^+^nuclei). Enteric glia as well as smooth muscle cells show increased EdU incorporation after 24h Aβ1-42, curli or LPS stimulation (mean ± SD; ANOVA glial nuclei p<0.005 and all nuclei p<0.005; **p<0.005 in Dunnett with medium); **C.** Representative confocal microscopy images and quantification of 72h medium-and curli-treated cultures, stained for γH2AX as a marker of DNA damage. γH2AX spot occupancy was quantified in glial nuclei (SOX10 segmentation; arrowhead indicates a γH2AX^+^/SOX10^+^ nucleus) and in all nuclei (based on DAPI segmentation), showing that glial as well as smooth muscle cells show amyloid-and LPS-induced double stranded DNA breaks (mean ± SD; ANOVA glial nuclei p<0.005 and all nuclei p<0.005; *p<0.05 and **p<0.005 in Dunnett with medium); **D.** Representative confocal microscopy images and quantification of 24h medium-and curli-treated cultures, stained for CellMask, SOD2 and the glia marker GFAP. Quantification of SOD2 intensity in the CellMask positive area reveals an increase after Aβ_1-42_, curli and LPS treatment compared to medium or Aβ_scr_ (mean ± SD; ANOVA p<0.005; **p<0.005 in Dunnett with medium). All scale bars 20 μm.

### In vivo injected amyloids induce oxidative stress and apoptosis in myenteric neurons and DNA damage in the smooth muscle layer

To further explore the oxidative stress status and DNA damage accumulation *in vivo,* we injected fluorescently labeled amyloids in the proximal colon wall of live mice and prepared myenteric whole mounts from the region around the injection site (**Fig. 3A**). While curli fibrils displayed a random distribution without apparent cellular uptake (**Suppl. Fig. 3A**), we found that injected Aβ_1-42_ accumulated in nuclei of myenteric neurons (**Suppl. Fig. 3B**). This phenomenon was not observed for its scrambled control (**Suppl. Fig. 3C, D**), and could be modeled *in vitro* by inducing membrane permeabilization in neurosphere-derived neurons but not by increasing the amyloid concentration or incubation time (**Suppl. Fig. 3E**), suggesting that oligomeric Aβ_1-42_ sticks to DNA of degenerating neurons.

**Figure 3.**
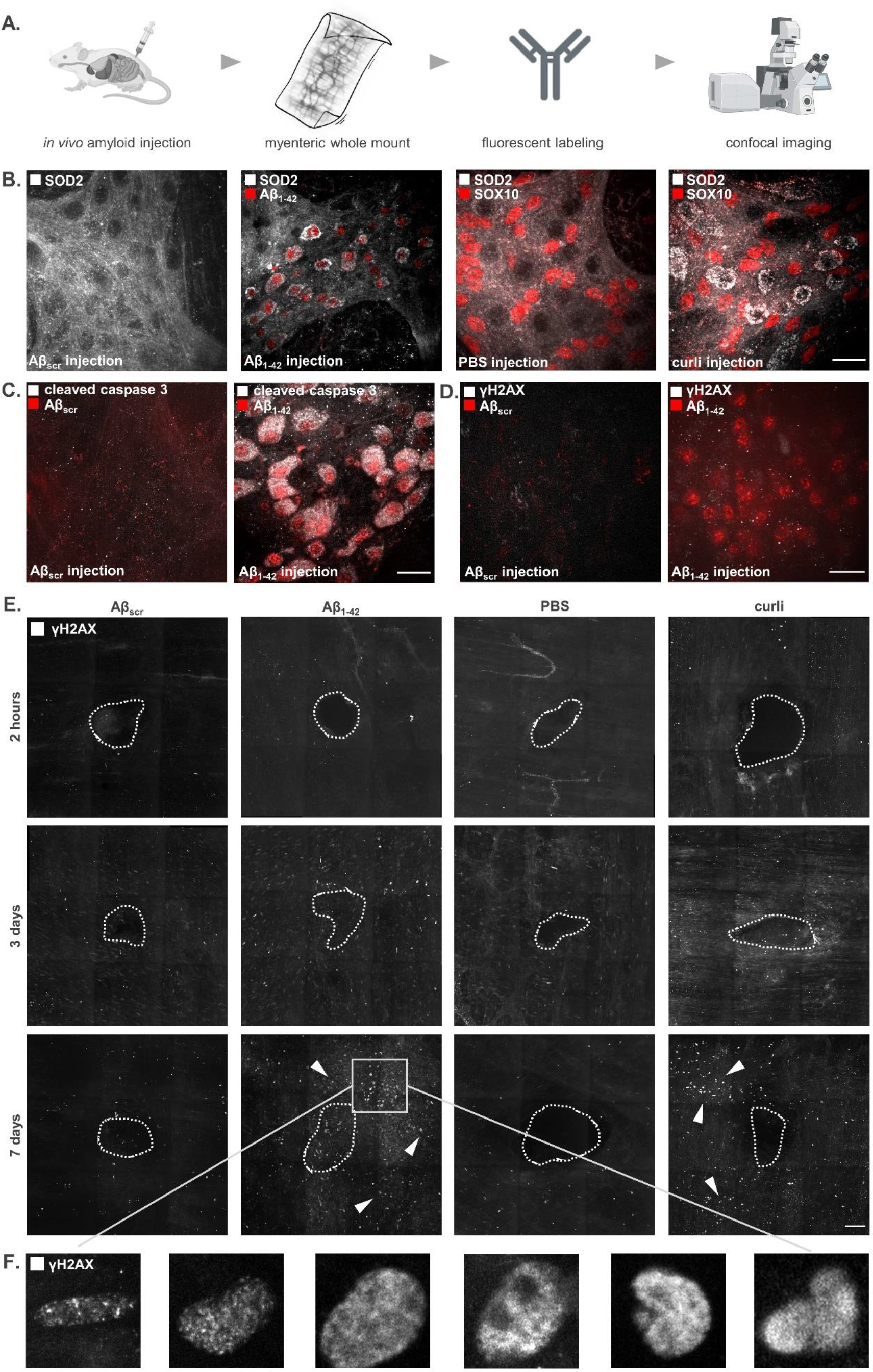
In vivo injected amyloids induce oxidative stress and apoptosis in myenteric neurons and DNA damage in the muscularis. **A.** Experimental workflow. Mice received an intramural injection Aβ_scr_-FAM, Aβ_1-42_-hilyte555, PBS or curli in the proximal colon and immunostainings were carried out on whole mount preparations from the region adjacent to the injection sites; **B.** Increased SOD2 levels 2h after the injection in Aβ_1-42_-positive neurons, which were absent in Aβ_scr_-injected mice. Similar upregulation was also observed after curli but not PBS injection; **C.** Cleaved caspase 3, a marker of apoptosis, was elevated in neurons that accumulate Aβ_1-42_ in their nuclei while it was absent in Aβ_scr_^-^ injected tissue, 2h post injection; **D.** Overt double-stranded DNA damage (γH2AX signal) was not detected in amyloid-bearing myenteric neurons at the 2h timepoint; **E.** γH2AX staining revealed a time-dependent accumulation of DNA damage (arrowheads) near the injection site (dashed line) upon Aβ_1-42_ or curli exposure but not after Aβ_scr_ and PBS injection; **F.** Zoomed and cropped images of nuclei with punctate or pan-nuclear γH2AX signal and altered morphology, reminiscent of different stages of cell death; scale bars **B-D** 20 μm and **E** 100 μm.

In line with RNA Seq and IF data, we observed marked SOD2 upregulation, which in this setting localized to myenteric neurons in curli-injected tissue and to neurons with nuclear Aβ_1-42_ (**Fig. 3B**). Increased SOD2 levels were not observed in Aβ_scr_- or PBS-injected animals, suggesting an amyloid-driven pathogenic process. Neurons with Aβ_1-42_^+^ nuclei had increased levels of cleaved caspase 3 (**Fig. 3C**), underscoring the capacity of amyloid to induce apoptosis in myenteric neurons. While these specific neurons did not show overt DNA damage (**Fig. 3D**), we did observe γH2AX accumulation in the region adjacent to an injection site, which could be identified based on a local disruption of the neuronal network (β_III_-tubulin), an accumulation of amyloid, and, depending on the time point, a cell infiltrate near the damaged part (DAPI) (**Suppl. Fig. 4A**). Accumulation of γH2AX-positive nuclei was induced by Aβ_1-42_ and curli but not by the control injections (Aβ_scr_ and PBS). This occurred in a time-dependent manner, with the largest difference between amyloid and control at the latest investigated time point, *i.e.,* 7 days after the injection (**Fig. 3E** arrowheads; the dashed line indicates the injection site). At this stage, nuclei showed punctate or more diffuse pan-nuclear γH2AX patterns and morphological deformations suggesting cell death (**Fig. 3F**). Inspection of confocal Z-planes showed that these nuclei were not abundant in the myenteric plexus but were found mainly in the muscle layer (**Suppl. Fig. 4B**; arrowheads indicate the myenteric plexus), which was consistent with the previously observed γH2AX accumulation in neurosphere-derived smooth muscle cells (**Fig. 2C**). Thus, amyloids can induce oxidative stress and DNA damage *in vivo* as well, which may result in neuronal cell death.

### Enteric glia initiate a pro-inflammatory response upon amyloid challenge

Neurosphere-derived glia displayed several features of an activated cell state (**Fig. 2**). In line with this, several of the upregulated genes pertained to an innate immune response (**Fig. 4A**). Therefore, we decided to further investigate the inflammatory response at the protein level by measuring pro-inflammatory cytokine concentrations in the cell culture supernatant of amyloid-treated myenteric networks. Curli induced the release of the pyrogen IL1β, and the chemokines CXCL1, CXCL2, CCL2, CCL3 and CCL5 (**Fig. 4B**). Even though similar trends were observed after Aβ_1-42_ stimulation, it only reached statistical significance for CXCL2, suggesting that curli is more immunogenic, at least at the tested concentration. To determine the cellular origin of the pro-inflammatory cytokines, we differentiated primary enteric neurospheres to glial cell cultures, or to enteric neurons (**Fig. 4C**). While the glial cultures contained a mixture of enteric glia and smooth muscle cells (**Suppl. Fig. 2A**), the neuronal cultures were enriched in enteric neurons but also contained a lower amount of glia and smooth muscle cells (**Suppl. Fig. 2B**). qPCR analyses showed that the response to Aβ_1-42_ and curli was more pronounced in enteric glia than in neurons, whereby curli induced cytokine expression with nearly the same potency as LPS (**Fig. 4C**). To confirm the cytokine-producing cell type *in vivo*, we performed fluorescence *in situ* hybridization for *Cxcl2* mRNA on whole mount preparations of Aβ_scr_ or Aβ_1-42_-injected colon, sacrificed 2h after injection (**Fig. 4D**). *Cxcl2* transcripts were enriched in enteric glia of the myenteric plexus (as counterstained with the nuclear marker SOX10) and significantly more abundant in enteric ganglia that contained neurons with nuclear Aβ_1-42_. In the same whole mounts, ganglia devoid of Aβ_1-42_-filled neurons were indistinguishable from Aβ_scr_-injected tissue (**Fig. 4D**). The spatial correlation of glial *Cxcl2* upregulation with nuclear Aβ_1-42_ accumulation in the neurons suggests an intricate neuro-immune interplay in amyloid-associated neurodegeneration. Since many of the observed chemokines are known to activate the adaptive immune system, we performed flow cytometry on curli-injected colon to measure peripheral immune cell influx. We observed elevated CD45^+^ immune cells counts in the muscularis 7 days after an intramural injection. At this time point, the CD45^+^CD3^+^TCRβ^+^ T-cell population specifically showed a significant increase, where curli exceeded that of sole PBS injection (**Fig. 4E**). Collectively, these data show that amyloids induce a local inflammatory environment near the ENS that is at least partly initiated by enteric glia and exacerbated by peripheral immune cell infiltration.

**Figure 4.**
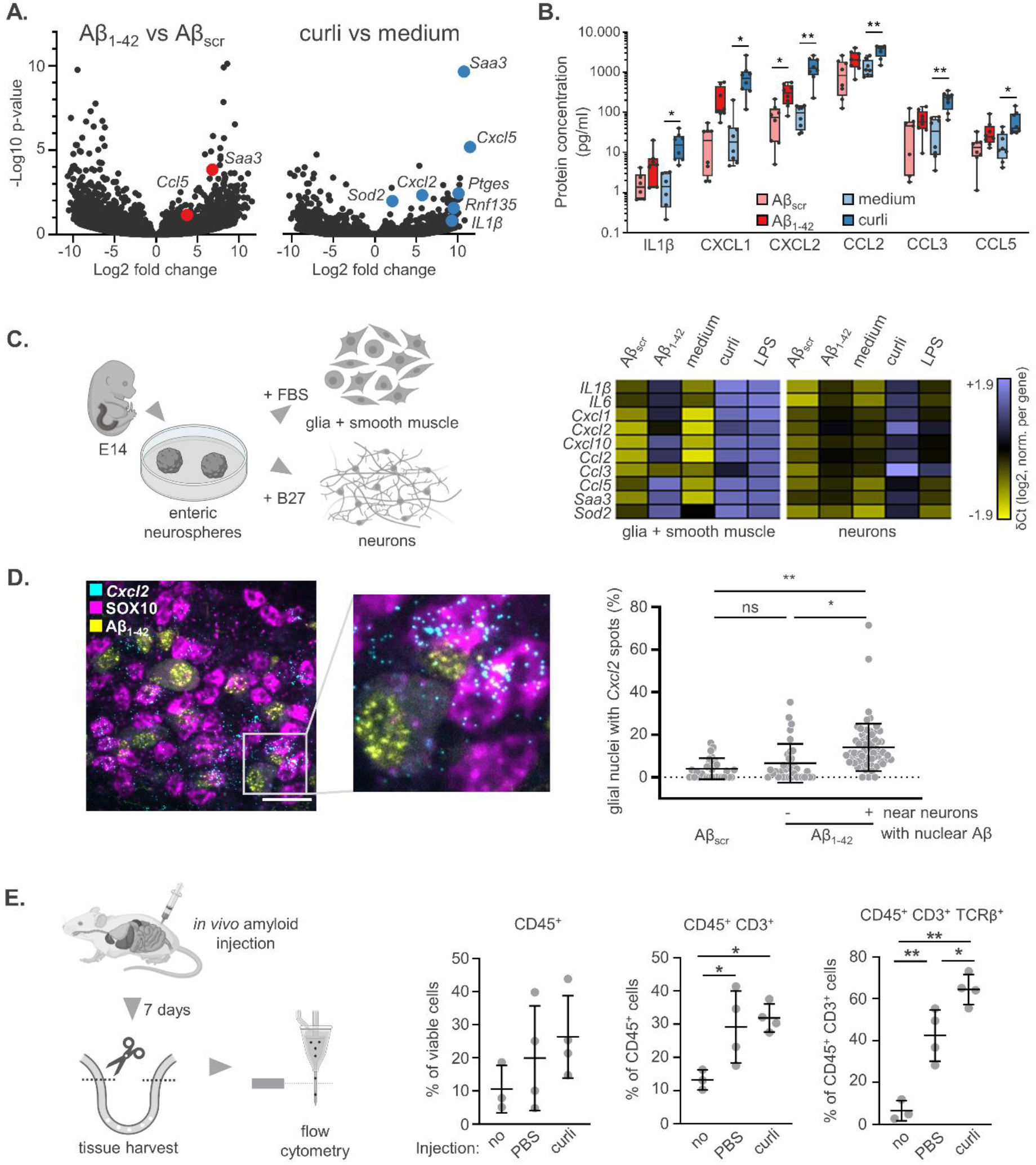
Enteric glia initiate a pro-inflammatory response upon amyloid challenge. **A.** Volcano plots of the RNA-Seq experiment described in **Fig. 1** show that many upregulated transcripts pertain to a pro-inflammatory response, whereby curli is more immunogenic than Aβ1- 42; **B.** Concentrations of pro-inflammatory cyto-and chemokines released into the culture medium, 24h after challenging myenteric networks with Aβ_1-42_ (vs Aβ_scr_) or curli (vs culture medium; n=8; glm with analyte/treatment/analyte*treatment all p<0.0001; *p<0.05 **p<0.005 in Holm-Sidak-corrected t-test); **C.** Neurosphere-derived enteric neurons and mixed glia/smooth muscle cell cultures cultures were challenged with amyloids or LPS for 24h. qPCR analysis revealed that mRNA encoding pro-inflammatory cytokines was induced mainly in the glial/smooth muscle cell cultures (n=3 independent cultures of each type); **D.** Fluorescence *in situ* hybridization for *Cxcl2*transcripts in myenteric whole mounts prepared from Aβ_1-42_-injected colon (2h post injection). The percentage of SOX10-counterstained glial nuclei that contains *Cxcl2* spots is increased in myenteric ganglia with Aβ_1-42_-bearing neurons. Ganglia with intact myenteric neurons (no nuclear Aβ_1-42_) in the same whole mounts are indistinguishable from Aβ_scr_-injected tissue (mean ± SD; n=3 animals with ≥10 images/animal and on average 35 SOX10^+^ nuclei/image; One-way ANOVA p<0.0001; *p<0.05 **p<0.005 in Tukey’s post-hoc). Scale bar 20 μm; **E.** Sterile PBS or curli were injected in the proximal colon of live mice, and muscularis tissue of the colon was processed for flow cytometry 7 days later. A control group that received no injection/operation was included as well (mean ± SD; n=4). The injected animals show a trend towards higher CD45^+^ immune cell influx. A population of T-cells (CD45^+^ CD3^+^ TCRβ^+^) was specifically enriched after curli compared to PBS injection (ANOVA CD45^+^ p=0.4025; CD45^+^ CD3^+^ p=0.0224; CD45^+^ CD3^+^TCRβ^+^ p=0.0001; *p<0.05 **p<0.005 in Tukey’s post-hoc).

## Discussion

During aging and inflammation, the ENS becomes exposed to human as well as bacterial amyloids^1,4^. This study shows that these amyloids are not innocent bystanders but activate pathogenic pathways that sustain local pathology and potentially also influence central neurodegeneration.

We observed a pronounced dysregulation of the cell cycle 24h after challenge with either amyloid type. The bulk RNA-Seq data primarily indicated cell cycle arrest but also contained a small hub pointing to smooth muscle cell proliferation. Since the subsequent *in vitro* experiments exposed a clear proliferative state of glial and smooth muscle cells, and given the difference in cellular composition of myenteric networks and neurosphere-derived glial cultures (which do not contain neurons) we believe that the myenteric neurons may be responsible for the apparent cell cycle arrest in the RNA-Seq dataset. Even though they are considered post-mitotic, they express cell cycle regulators, albeit serving alternate functions such as neurite morphogenesis and synaptic plasticity^33^. Also, in central neurons, Aβ has been shown to induce ectopic cell cycle re-entry, which may be detrimental^34–36^ but was recently also shown to protect against Aβ-induced apoptosis^37^.

The observed proliferative state of enteric glia was accompanied by an oxidative stress response (evident by SOD2 upregulation) and accumulation of DNA damage (nuclear γH2AX spot occupancy). Consistent with this, amyloids have been shown to promote genomic instability by increasing oxidative DNA damage in the CNS^32^. They also reduce DNA repair capacity and sensitize cells to otherwise nonlethal oxidative injury^38^. While a DNA damage response was only evident in non-neuronal (cycling) cells in our relatively short-term *in vitro* and injection models, other literature reports have described increased γH2AX signal in neurons of AD patients as well^22^. In all employed models (primary myenteric networks, neurosphere-derived glial and neuronal cultures, and *in vivo* injections) we consistently found SOD2 upregulation after challenge with either amyloid type. This reactive oxygen species (ROS) scavenging enzyme is localized in the mitochondrial matrix and represents the first line of defense against oxidative stress. SOD2 can be induced by oxidative stress and TLR2 signaling, and its enzymatic activity is modulated by cyclin B1, cyclin-dependent kinase 1 and p53^39,40^. The current RNA-Seq data show that all 3 regulatory genes were downregulated after amyloid stimulation, suggesting dysregulation of the adaptive oxidative stress response. We suspect that the faster proliferation rate in combination with a dysregulated oxidative stress response may have rendered glial and smooth muscle cells more vulnerable to DNA damage. In the *in vivo* injection model, higher SOD2 levels were only evident in degenerating Aβ_1-42_^+^ neurons. This may be the result of local high concentrations and mechanical stress in this model but shows that this response is preserved among different cell types.

We found that amyloids, and in particular curli, are potent inducers of an immune response in the ENS. While both amyloid types show no homology in their amino acid sequences, they are structurally related in the sense that they form β-sheets which are recognized by TLRs^12,41–43^. The enhanced immunogenicity of fibrillar curli compared to oligomeric Aβ_1-42_ may be explained by formulation differences, since curli fibers contain nucleic acids that represent an additional TLR substrate compared to the more pure Aβ_1-42_ oligomers^44^. Many of the cytokines that we identified in the ENS have a clear link with amyloid-induced neuropathology in the brain as well. IL1β acts as a pro-inflammatory mediator and pyrogen that is released after intracellular activation of the NLRP3 inflammasome, for which Aβ and curli are known inducers^45,46^. Blood levels of IL1β, NLRP3 and CXCL2 have been positively correlated with the abundance of curli-producing bacteria in stool, as measured in healthy controls, cognitively impaired patients with and without amyloid pathology in the brain^47^. CXCR2, the receptor for CXCL1 and CXCL2, is expressed by microglia and its abundance was increased after intrahippocampal injection of Aβ_1-42_, while a specific antagonist inhibited microgliosis, oxidative stress and neuronal loss^48^. Intracerebroventricular injection of Aβ1-40 in WT mice induced the expression of CCL3 and CCR5, the receptor for CCL3/4/5, followed by astro-and microgliosis in the hippocampus^49^. Aβ1-40 injection in CCL3-/- or CCR5-/- mouse brains attenuated astro-and microgliosis, synaptic dysfunction and cognitive defects, suggesting that the CCL3/CCR5 pathway mediates neuroinflammation and therefore contributes to neurodegeneration. CCL2 is expressed by central neurons and astrocytes and its levels in CSF at baseline correlate with a faster cognitive decline and could even be used as a biomarker in combination with CSF Tau, pTau and Aβ_1-42_ to predict future conversion to AD^50,51^. We now show that the same cytokines are involved in the innate immune response towards amyloids in the GI tract. Another striking parallel between the CNS and our data obtained in the ENS was recently provided by a study where rat hippocampal astrocytes were exposed to Aβ1-40. The amyloid-treated astrocytes showed increased proliferation, inflammatory cytokine and SOD2 upregulation, as well as higher ROS levels^52^. This shows that not only the cytokine palette but also the activated glial state represent significant parallel manifestations in the GI tract and the brain. If and to what extent these pathways are involved in reciprocal gut-brain communication and pathology transfer remains to be determined.

The current study shows the pathogenic potential of human and bacterial amyloids in the GI tract. While the long-term consequences on GI homeostasis and contribution to CNS pathology should be studied more in depth, this insight may open novel avenues for novel targets for early diagnosis and therapeutic intervention of AD and related amyloid pathologies.

## Acknowledgements

This work was funded by The Research Foundation Flanders (FWO project G017618N to JPT and WDV, FWO I003420N to WDV and FWO IRI I000321N to WDV) and the University of Antwerp (TOP-BOF project 35020 to JPT and WDV and BOF-KP 39616 to PV). CT is supported by NIH grants AI153325, AI151893, and AI148770. We thank dr. Esther Bartholomeus of the Center for Medical Genetics, University of Antwerp, for technical support during RNA sequencing.

## Author contributions

PV, JPT, WDV designed the research; PV, SVR, NDL, RV, AC, MG, SIM performed the research; PV, GG, SIM analyzed the data; CT, KHS contributed new reagents/analytical tools; PV, WDV wrote the paper. All authors critically revised and approved the manuscript.

## Supplementary figures

**Supplementary figure 1.**
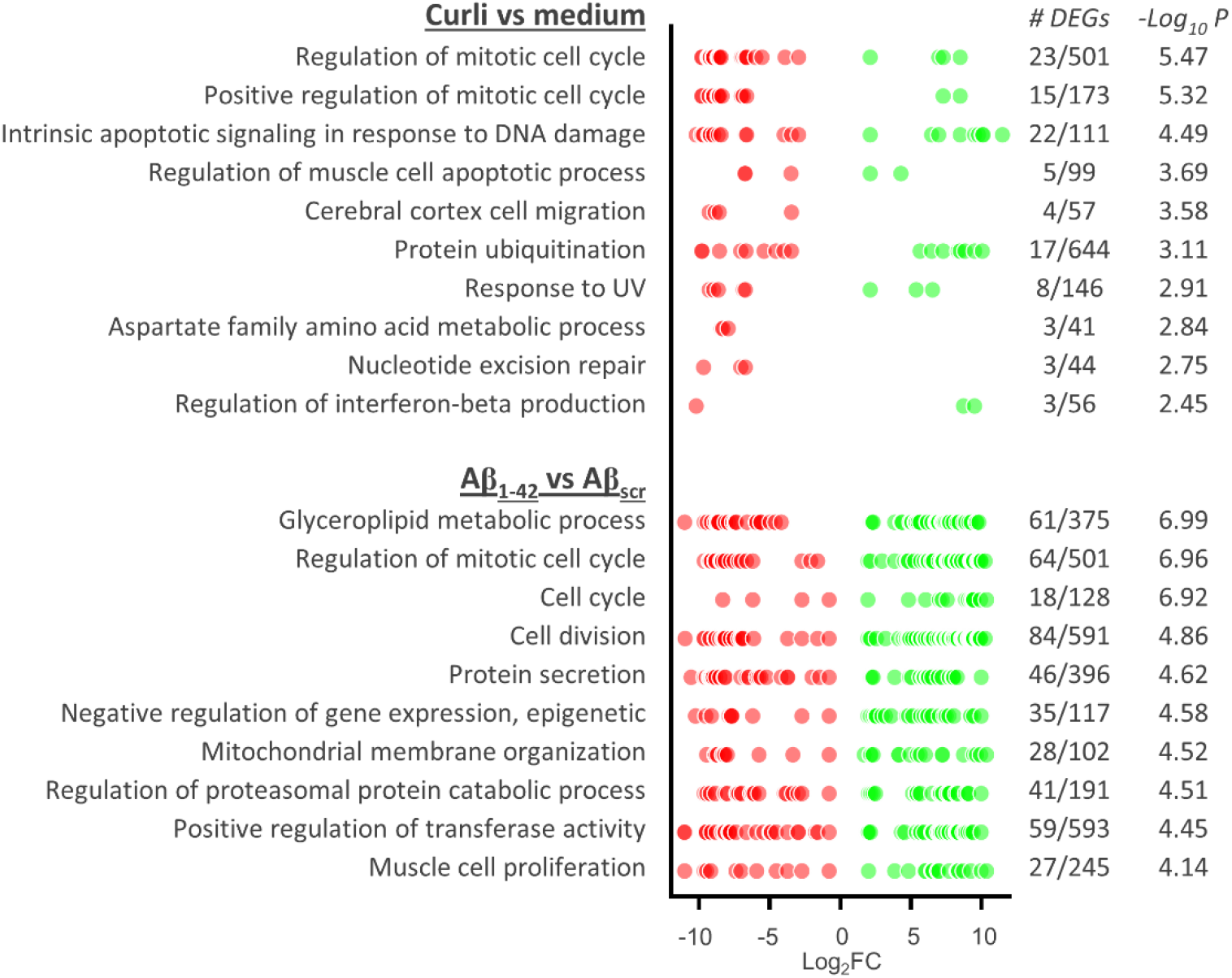
Enriched Gene Ontology terms 24h after amyloid challenge. Top 10 enriched terms for each amyloid type, with the Log2 Fold Change of DEGs shown in red (downregulated) and green (upregulated). The number of DEGs and p-value of term enrichment are reported as well.

**Supplementary figure 2.**
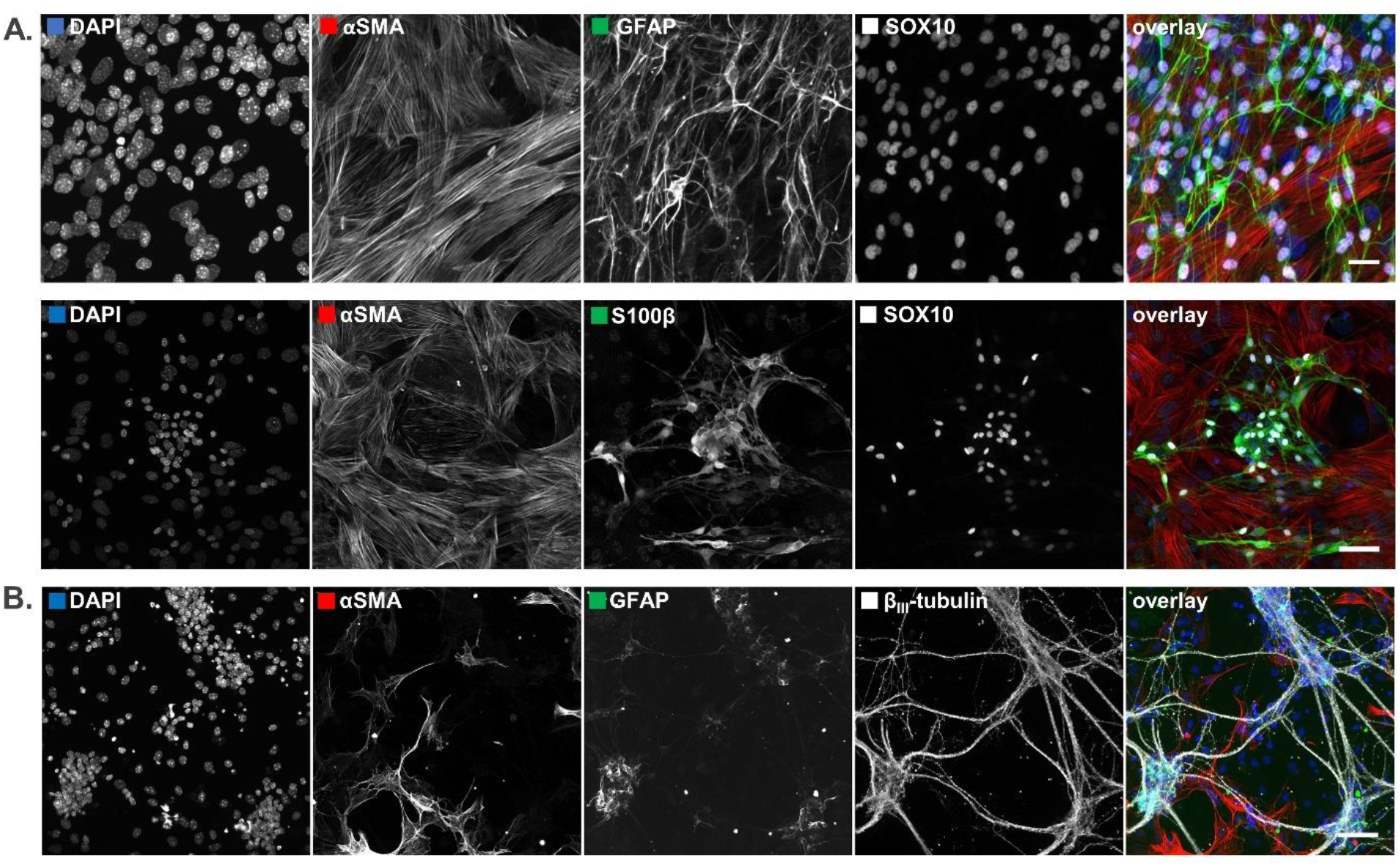
Cell types present in enteric neurosphere-derived cultures. Neurosphere-derived cultures were differentiated to enriched glia or neuronal cell cultures. **A.** Confocal microscopy image show that neurosphere-derived glial cultures are mainly composed of enteric glia (GFAP, S100β and SOX10) and smooth muscle cells (αSMA); **B.** A differentiated neuronal culture stained for the neuron marker β_III_-tubulin, and for the glial and smooth muscle markers GFAP and αSMA, respectively. In addition to an extensive neuronal network, these cultures contain a low number of glia and smooth muscle cells. All scale bars 50 μm.

**Supplementary figure 3.**
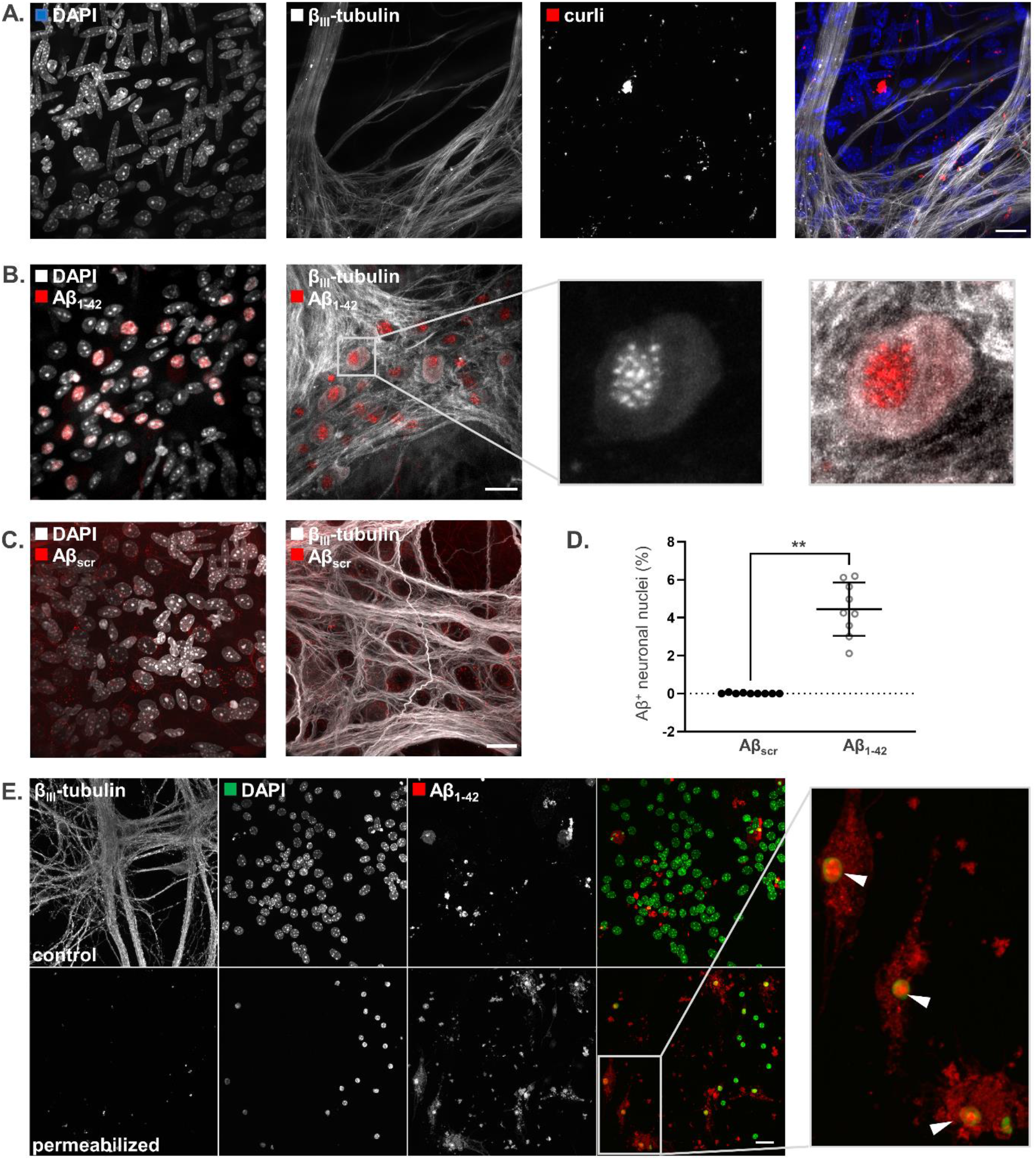
Injected amyloids accumulate differently in the myenteric plexus. Fluorescently labeled curli, Aβ_1-42_ or Aβ_scr_ were injected into the proximal colon wall of live WT mice, which were sacrificed 2h later. Then, whole mounts were prepared from the region close to the injection sites to study amyloid distribution and uptake near the myenteric plexus; **A.** Injected curli showed a stochastic distribution pattern without apparent cellular uptake; **B.** Aβ_1-42_ accumulated in nuclei of myenteric neurons, as shown by DAPI and β_III_-tubulin counterstaining; **C.** Injected Aβ_scr_ did not accumulate in neuronal nuclei; **D.** Quantification revealed that the percentage of neuronal nuclei with amyloid signal was significantly higher for Aβ_1-42_ than Aβ_scr_ (n=3 animals * 3 whole mounts; p<0.005 in t-test); **E.** Neurosphere-derived enteric neurons were exposed to 1 μM Aβ_1-42_-hilyte555 in the presence or absence of 0.02% Triton X-100 for 24h. Amyloid accumulation in nuclei was observed in permeabilized cells but not in control cultures. All scale bars 20 μm.

**Supplementary figure 4.**
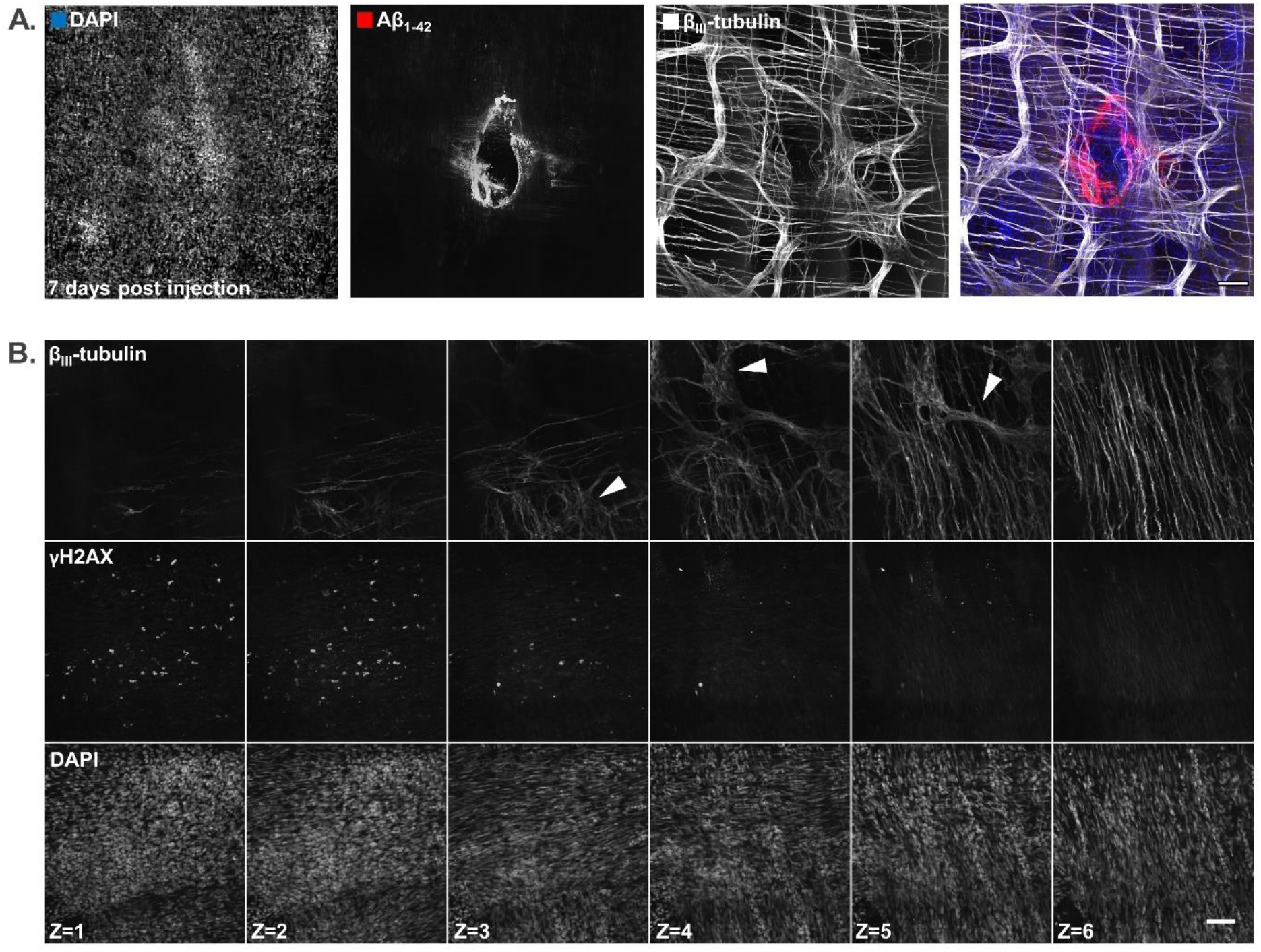
Identification of an injection site and localization of γH2AX^+^ nuclei to the muscle layer. **A.** Identification of an injection site based on an accumulation of amyloid, a local disruption of the neuronal network (β_III_-tubulin), and a cell infiltrate near the damaged part (DAPI; 7 days post injection); **B.** Confocal images taken at different Z-positions indicate that the majority of γH2AX^+^nuclei are localized in the muscle layer and not in ganglia of the myenteric plexus (myenteric plexus is indicated with arrowheads). All scale bars 100 μm.

## References

1. Cryan JF, O’Riordan KJ, Sandhu K, Peterson V, Dinan TG. The gut microbiome in neurological disorders. Lancet Neurol. 2020;19(2):179–194.

2. Kim GH, Lee YC, Kim TJ, et al. Risk of neurodegenerative diseases in patients with inflammatory bowel disease: a nationwide population-based cohort study. J Crohns Colitis. 2022;16(3):436–443.

3. Zhang B, Wang HE, Bai Y-M, et al. Inflammatory bowel disease is associated with higher dementia risk: a nationwide longitudinal study. Gut. 2021;70(1):85.

4. Manocha GD, Floden AM, Miller NM, et al. Temporal progression of Alzheimer’s disease in brains and intestines of transgenic mice. Neurobiol Aging. 2019;81:166–176.

5. Semar S, Klotz M, Letiembre M, et al. Changes of the enteric nervous system in amyloid-β protein precursor transgenic mice correlate with disease progression. J Alzheimer’s Dis. 2013;36(1):7–20.

6. Sun Y, Sommerville NR, Liu JYH, et al. Intra-gastrointestinal amyloid-beta1-42 oligomers perturb enteric function and induce Alzheimer’s disease pathology. J Physiol. 2020;598(19):4209–4223.

7. Westwell-Roper C, Verchere CB. Modulation of Innate Immunity by Amyloidogenic Peptides. Trends Immunol. 2019;40(8):762–780.

8. Tytgat HLP, Nobrega FL, van der Oost J, de Vos WM. Bowel biofilms: tipping points between a healthy and compromised gut? Trends Microbiol. 2019;27(1):17–25.

9. Sobieszczanska B, Pawlowska B, Duda-Madej A, et al. Effect of amyloid curli fibrils and curli CsgA monomers from Escherichia coli on in vitro model of intestinal epithelial barrier stimulated with cytokines. Int J Med Microbiol. 2019;309(5):274–282.

10. Arai H, Lee VM, Messinger ML, Greenberg BD, Lowery DE, Trojanowski JQ. Expression patterns of beta-amyloid precursor protein (beta-APP) in neural and nonneural human tissues from Alzheimer’s disease and control subjects. Ann Neurol. 1991;30(5):686–693.

11. Puig KL, Lutz BM, Urquhart SA, et al. Overexpression of mutant amyloid-beta protein precursor and presenilin 1 modulates enteric nervous system. J Alzheimer’s Dis. 2015;44(4):1263–1278.

12. Tükel C, Nishimori JH, Wilson RP, et al. Toll-like receptors 1 and 2 cooperatively mediate immune responses to curli, a common amyloid from enterobacterial biofilms. Cell. Microbiol. 2010;12(10):1495–1505.

13. Friedland RP, McMillan JD, Kurlawala Z. What are the molecular mechanisms by which functional bacterial amyloids influence amyloid beta deposition and neuroinflammation in neurodegenerative disorders? Int J MolSci. 2020;21(5):1652.

14. Bhoite SS, Han Y, Ruotolo BT, Chapman MR. Mechanistic insights into accelerated α-synuclein aggregation mediated by human microbiome-associated functional amyloids. J Biol Chem. 2022;298(7):102088.

15. Wang C, Lau CY, Ma F, Zheng C. Genome-wide screen identifies curli amyloid fibril as a bacterial component promoting host neurodegeneration. PNAS. 2021;118(34).

16. Chen SG, Stribinskis V, Rane MJ, et al. Exposure to the functional bacterial amyloid protein curli enhances alpha-synuclein aggregation in aged fischer 344 rats and caenorhabditis elegans. Sci Rep. 2016;6(1):34477.

17. Sampson TR, Challis C, Jain N, et al. A gut bacterial amyloid promotes α-synuclein aggregation and motor impairment in mice. eLife. 2020;9:e53111.

18. Liu JYH, Sun MYY, Sommerville N, et al. Soy flavonoids prevent cognitive deficits induced by intra-gastrointestinal administration of beta-amyloid. Food Chem Toxicol. 2020;141:111396.

19. Guglielmotto M, Giliberto L, Tamagno E, Tabaton M. Oxidative stress mediates the pathogenic effect of different Alzheimer’s disease risk factors. Front Aging Neurosci. 2010;:2.

20. Leng F, Edison P. Neuroinflammation and microglial activation in Alzheimer disease: where do we go from here? Nat Rev Neurol. 2021;17(3):157–172.

21. Sanders OD, Rajagopal L, Rajagopal JA. The oxidatively damaged DNA and amyloid-β oligomer hypothesis of Alzheimer’s disease. Free Radic Biol Med. 2022;179:403–412.

22. Shanbhag NM, Evans MD, Mao W, et al. Early neuronal accumulation of DNA double strand breaks in Alzheimer’s disease. Acta Neuropathol Commun. 2019;7(1):77.

23. Miller AL, Bessho S, Grando K, Tukel C. Microbiome or infections: amyloid-containing biofilms as a trigger for complex human diseases. Front Immunol. 2021;12:638867.

24. Nicastro LK, Tursi SA, Le LS, et al. Cytotoxic curli intermediates form during Salmonella biofilm development. J Bacteriol. 2019;201(18).

25. Tükel C, Raffatellu M, Humphries AD, et al. CsgA is a pathogen-associated molecular pattern of Salmonella enterica serotype Typhimurium that is recognized by Toll-like receptor 2. Mol Microbiol. 2005;58(1):289–304.

26. Grundmann D, Klotz M, Rabe H, Glanemann M, Schafer KH. Isolation of high-purity myenteric plexus from adult human and mouse gastrointestinal tract. Sci Rep. 2015;5:9226.

27. Zhou Y, Zhou B, Pache L, et al. Metascape provides a biologist-oriented resource for the analysis of systems-level datasets. Nat Commun. 2019;10(1):1523.

28. Schindelin J, Rueden CT, Hiner MC, Eliceiri KW. The ImageJ ecosystem: An open platform for biomedical image analysis. Mol Reprod Dev. 2015;82(7-8):518–529.

29. Sturn A, Quackenbush J, Trajanoski Z. Genesis: cluster analysis of microarray data. Bioinformatics. 2002;18(1):207–208.

30. Gupta S, Goswami P, Biswas J, et al. 6-Hydroxydopamine and lipopolysaccharides induced DNA damage in astrocytes: involvement of nitric oxide and mitochondria. Mutat Res Genet Toxicol Environmenl Mutagen. 2015;778:22–36.

31. Zhang K, Wu S, Li Z, Zhou J. MicroRNA-211/BDNF axis regulates LPS-induced proliferation of normal human astrocyte through PI3K/AKT pathway. Biosci Rep. 2017;37(4).

32. Mao P, Reddy PH. Aging and amyloid beta-induced oxidative DNA damage and mitochondrial dysfunction in Alzheimer’s disease: implications for early intervention and therapeutics. Biochim Biophys Acta. 2011;1812(11):1359–1370.

33. Frank CL, Tsai L-H. Alternative functions of core cell cycle regulators in neuronal migration, neuronal maturation, and synaptic plasticity. Neuron. 2009;62(3):312–326.

34. Kodis EJ, Choi S, Swanson E, Ferreira G, Bloom GS. N-methyl-D-aspartate receptor-mediated calcium influx connects amyloid-beta oligomers to ectopic neuronal cell cycle reentry in Alzheimer’s disease. Alzheimers Dement. 2018;14(10):1302–1312.

35. Lopes JP, Oliveira CR, Agostinho P. Cdk5 acts as a mediator of neuronal cell cycle re-entry triggered by amyloid-beta and prion peptides. Cell Cycle. 2009;8(1):97–104.

36. Seward ME, Swanson E, Norambuena A, et al. Amyloid-beta signals through tau to drive ectopic neuronal cell cycle re-entry in Alzheimer’s disease. J Cell Science. 2013;126(Pt 5):1278–1286.

37. Ippati S, Deng Y, van der Hoven J, et al. Rapid initiation of cell cycle reentry processes protects neurons from amyloid-beta toxicity. PNAS 2021;118(12).

38. Cardinale A, Racaniello M, Saladini S, et al. Sublethal doses of beta-amyloid peptide abrogate DNA-dependent protein kinase activity. J Biol Chem. 2012;287(4):2618–2631.

39. Liu YD, Yu L, Ying L, et al. Toll-like receptor 2 regulates metabolic reprogramming in gastric cancer via superoxide dismutase 2. Int j Cancer. 2019;144(12):3056–3069.

40. Candas D, Li JJ. MnSOD in oxidative stress response-potential regulation via mitochondrial protein influx. Antiox Redox Signal. 2014;20(10):1599–1617.

41. Hughes C, Choi ML, Yi J-H, et al. Beta amyloid aggregates induce sensitised TLR4 signalling causing long-term potentiation deficit and rat neuronal cell death. Comm Biol. 2020;3(1):79.

42. Richard KL, Filali M, Préfontaine P, Rivest S. Toll-like receptor 2 acts as a natural innate immune receptor to clear amyloid β1-42 and delay the cognitive decline in a mouse model of Alzheimer’s disease. J Neurosci. 2008;28(22):5784–5793.

43. Tükel C, Wilson RP, Nishimori JH, Pezeshki M, Chromy BA, Bäumler AJ. Responses to amyloids of microbial and host origin are mediated through toll-like receptor 2. Cell Host Microbe. 2009;6(1):45–53.

44. Tursi SA, Lee EY, Medeiros NJ, et al. Bacterial amyloid curli acts as a carrier for DNA to elicit an autoimmune response via TLR2 and TLR9. PLoSpathogens. 2017;13(4):e1006315–e1006315.

45. Halle A, Hornung V, Petzold GC, et al. The NALP3 inflammasome is involved in the innate immune response to amyloid-beta. Nat Immunol. 2008;9(8):857–865.

46. Rapsinski GJ, Wynosky-Dolfi MA, Oppong GO, et al. Toll-like receptor 2 and NLRP3 cooperate to recognize a functional bacterial amyloid, curli. Infect Immun. 2015;83(2):693.

47. Cattaneo A, Cattane N, Galluzzi S, et al. Association of brain amyloidosis with pro-inflammatory gut bacterial taxa and peripheral inflammation markers in cognitively impaired elderly. Neurobiol Aging. 2017;49:60–68.

48. Ryu JK, Cho T, Choi HB, Jantaratnotai N, McLarnon JG. Pharmacological antagonism of interleukin-8 receptor CXCR2 inhibits inflammatory reactivity and is neuroprotective in an animal model of Alzheimer’s disease. J Neuroinflam. 2015;12(1):144.

49. Passos GF, Figueiredo CP, Prediger RD, et al. Role of the macrophage inflammatory protein-1alpha/CC chemokine receptor 5 signaling pathway in the neuroinflammatory response and cognitive deficits induced by beta-amyloid peptide. Am J Pathol. 2009;175(4):1586–1597.

50. Sokolova A, Hill MD, Rahimi F, Warden LA, Halliday GM, Shepherd CE. Monocyte chemoattractant protein-1 plays a dominant role in the chronic inflammation observed in Alzheimer’s disease. Brain Pathol. 2009;19(3):392–398.

51. Westin K, Buchhave P, Nielsen H, Minthon L, Janciauskiene S, Hansson O. CCL2 is associated with a faster rate of cognitive decline during early stages of Alzheimer’s disease. PloS one. 2012;7(1):e30525.

52. Lennol MP, Canelles S, Guerra-Cantera S, et al. Amyloid-β1-40 differentially stimulates proliferation, activation of oxidative stress and inflammatory responses in male and female hippocampal astrocyte cultures. Mech Ageing Dev. 2021;195:111462.

